# Anatomical predictors of successful prism adaptation in chronic visual neglect

**DOI:** 10.1101/144956

**Authors:** Marine Lunven, Gilles Rode, Clémence Bourlon, Christophe Duret, Raffaella Migliaccio, Emmanuel Chevrillon, Michel Thiebaut de Schotten, Paolo Bartolomeo

## Abstract

Visual neglect is a frequent and disabling consequence of right hemisphere damage. Previous work demonstrated a probable role of posterior callosal dysfunction in the chronic persistence of neglect signs. Prism adaptation is a non-invasive and convenient technique to rehabilitate chronic visual neglect, but it is not effective in all patients. Here we aimed to assess the hypothesis that prism adaptation improves left neglect by facilitating compensation through the contribution of the left, undamaged hemisphere. We assessed the relationship between prism adaptation effects, cortical thickness and white matter integrity in a group of 14 patients with unilateral right-hemisphere strokes and chronic visual neglect. Results showed that patients who benefitted from prism adaptation had thicker cortex in temporo-parietal, prefrontal and cingulate areas of the left, undamaged hemisphere. Additionally, these patients had a higher fractional anisotropy value in the body and genu of the corpus callosum. Results from normal controls show that these callosal regions connect temporo-parietal, sensorimotor and prefrontal areas. Finally, shorter time intervals from the stroke tended to improve patients’ response to prism adaptation. We concluded that prism adaptation may improve left visual neglect by promoting the contribution of the left hemisphere to neglect compensation. These results support current hypotheses on the role of the healthy hemisphere in the compensation for stroke-induced, chronic neuropsychological deficits, and suggest that prism adaptation can foster this role by exploiting sensorimotor/prefrontal circuits, especially when applied at early stages post-stroke.

## 1. Introduction

A major challenge of precision medicine is to relate individual patients’ characteristics to their clinical outcomes. Patients with right hemisphere damage and left visual neglect are unable to pay attention to left-sided objects, with consequent loss of autonomy and poor functional outcome (Bartolomeo, 2014; Heilman & Van Den Abell, 1980; Mesulam, 1981). At least 80% of patients with a right hemispheric stroke show signs of visual neglect in the acute stage (Azouvi et al., 2002). More rarely, right-sided neglect can occur after left hemisphere strokes (Beis et al., 2004), but its manifestations are less severe and patients tend to show a better recovery. Impaired integration of attention-related processes within the right hemisphere (Corbetta & Shulman, 2011; Doricchi, Thiebaut de Schotten, Tomaiuolo, & Bartolomeo, 2008), as well as between the hemispheres (Bartolomeo, Thiebaut de Schotten, & Doricchi, 2007; Heilman & Adams, 2003), contributes to neglect behaviour. About half of neglect patients still show signs of neglect one year or more after their stroke (Lunven et al., 2015), perhaps because posterior callosal dysfunction prevents attention-critical regions in the healthy hemisphere to access information processed by the damaged right hemisphere (Lunven et al., 2015).

Prism adaptation (PA) is a convenient technique to improve the ratio of recovering patients, because it is totally non-invasive and easy to administer (Rossetti et al., 1998; Yang, Zhou, Chung, Li-Tsang, & Fong, 2013). Prismatic goggles displace the whole visual field rightwards and induce an initial rightward bias in manual reaching movements, which disappears after a few trials. After removal of the prisms, reaching initially deviates towards the left neglected side, and neglect eventually disappears. Unfortunately, neglect does not improve in all treated patients (Luauté et al., 2006; Rode et al., 2015; Rousseaux, Bernati, Saj, & Kozlowski, 2006; Saj, Cojan, Vocat, Luauté, & Vuilleumier, 2013), for unknown reasons (Barrett, Goedert, & Basso, 2012; Chokron, Dupierrix, Tabert, & Bartolomeo, 2007).

Hypotheses on the neural mechanisms of PA in healthy subjects suggested processes of strategic control, dependent on the posterior parietal cortex, and of spatial realignment, associated to the cerebellum (Pisella, Rode, Farnè, Tilikete, & Rossetti, 2006; Striemer & Danckert, 2010). Neuroimaging evidence in healthy individuals showed PA-induced modulation of activity in these regions (Chapman et al., 2010; Clower et al., 1996; Crottaz-Herbette, Fornari, & Clarke, 2014; Danckert, Ferber, & Goodale, 2008; Luauté et al., 2009), together with prefrontal regions including the cingulate cortex (Danckert et al., 2008). Martín-Arévalo et al. (2016) showed that leftward PA alters transcallosal motor inhibition between the two motor cortices in the healthy brain, thus suggesting the possibility that PA also modulates inter-hemispheric interactions. Also consistent with this possibility, a recent model (Clarke & Crottaz-Herbette, 2016) postulates that the inferior parietal lobule (IPL) in the left hemisphere relays left- and right-sided visual stimuli to the dorsal system bilaterally. PA-induced input from the left IPL to the right dorsal system would thus be likely to increase inhibition exerted by the right hemisphere on the left hemisphere, thereby reducing signs of left neglect.

However, several gaps remain in our knowledge of the mechanisms of action of PA in visual neglect (Barrett et al., 2012). Also, there is no agreement on the potential lesion sites in the right hemisphere that might predict PA failure. Potential candidates include the posterior parietal cortex and superior frontal gyrus (Rousseaux et al., 2006), the infero-posterior parietal cortex (Luauté et al., 2006), the intra-parietal region and white matter around the inferior parietal lobule and the middle frontal gyrus (Sarri et al., 2008), the occipital lobe (Serino, Angeli, Frassinetti, & Làdavas, 2006), the posterior occipito-parietal area and right cerebellum (patient 5 in Saj et al., 2013). PA-induced improvement of neglect correlated with modifications of BOLD responses or regional blood flow (as estimated by PET), suggesting increased activity in bilateral fronto-parietal networks (Saj et al., 2013), right cerebellum, left thalamus, left temporo-occipital cortex, and decreased activity in left medial temporal cortex and right posterior parietal cortex (Luauté et al., 2006). In brain-damaged patients without neglect, damage to posterior parietal cortex and cerebellum generally impairs adaptation to PA (Martin, Keating, Goodkin, Bastian, & Thach, 1996; Pisella et al., 2005; Weiner, Hallett, & Funkenstein, 1983)

Drawing on our previous evidence suggesting a role for posterior callosal dysfunction in the chronic persistence of neglect (Lunven et al., 2015), we hypothesized that PA might improve neglect by facilitating compensation through the contribution of the left, undamaged hemisphere. To test this hypothesis, we assessed the relationship between PA effects, cortical thickness and white matter integrity in a group of 14 patients with unilateral right-hemisphere stroke and chronic left visual neglect.

## 2. Methods

### 2.1 Standard Protocol Approvals, Registrations, and Patient Consents

The study was promoted by the Inserm (C10-48) and approved by the Ile-de-France I Ethics Committee (2011-juin-12642). All participants gave written informed consent to participate in the study. No part of the study procedures was pre-registered prior to the research being conducted.

### 2.2 Subjects

A group of 14 patients with a single right hemisphere haemorrhagic or ischemic stroke, which had occurred at least 3 months before testing, showing stable signs of left visual neglect, gave written informed consent to participate in the study. None of the patients had a prior history of neurological disease or psychiatric disorders. Table 1 reports patients’ demographic, clinical and neuropsychological data. Visual neglect was assessed by using six paper-and-pencil tasks: letter cancellation (Mesulam, 1985), copy of a linear drawing of a landscape (Gainotti, D’Erme, & de Bonis, 1989), drawing from memory of a daisy (Rode, Rossetti, & Boisson, 2001); text reading (Azouvi et al., 2006), bisection of five 20-cm horizontal lines (Azouvi et al., 2006) and landmark task (Harvey & Milner, 1999). All patients performed these tests on three occasions: (1) the day before PA, (2) two hours before PA, and (3) soon after PA. For each assessment, a neglect severity score was computed as previously reported (Lunven et al., 2015), by using the percentage of left omissions for letter cancellation, drawing and reading tasks, the percentage of rightwards deviation for line bisection, and the percentage of left lateralised errors for the landmark task. Ten age-matched healthy subjects (mean age, 62.8 years; SD, 4.54; Mann-Whitney-Wilcoxon test, p ns) served as controls for neuroimaging.

**Table 1:**
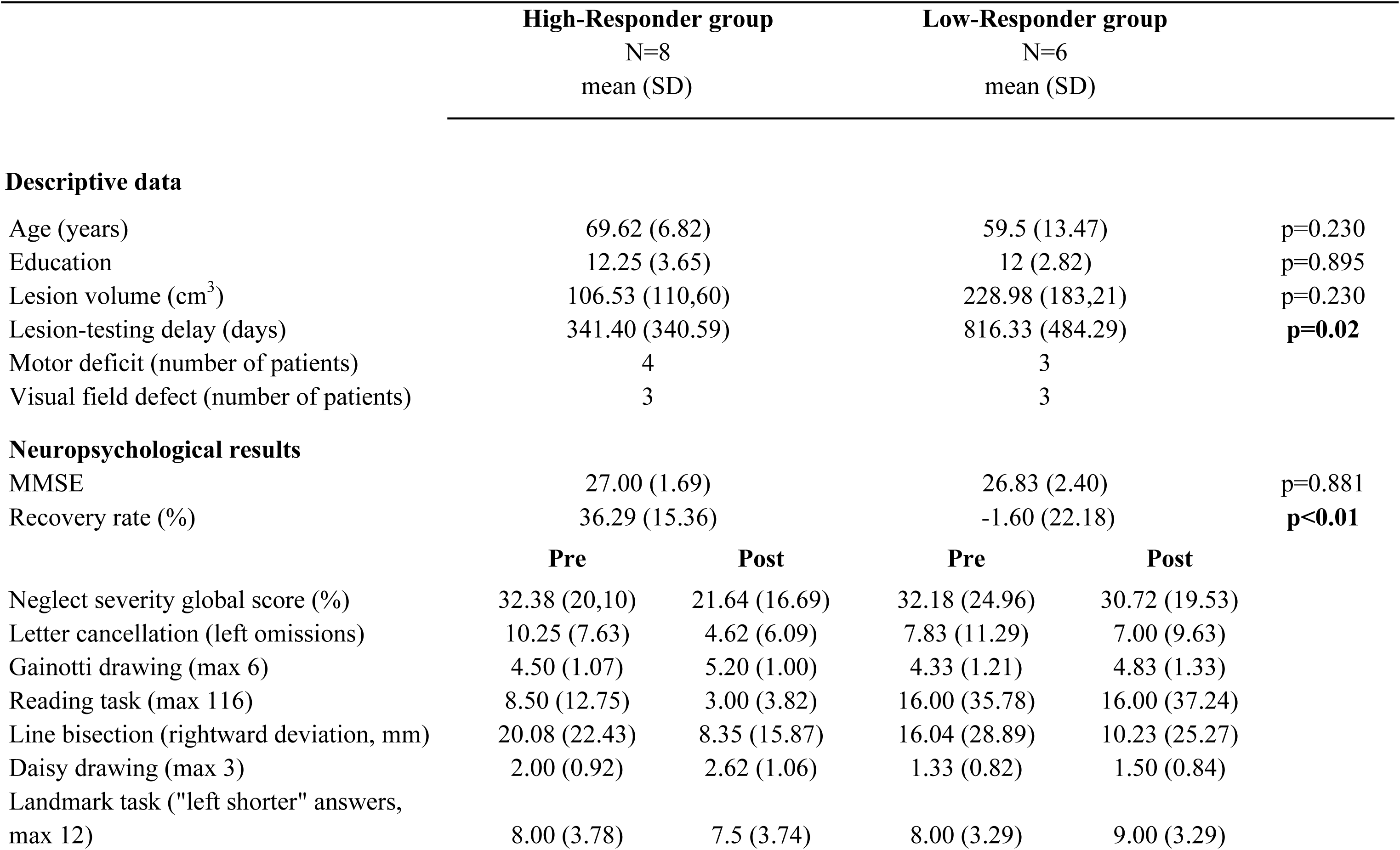
Demographical and clinical characteristics of patients, with their performance on visuospatial tests before and after prism adaptation.

| Descriptive data | High-Responder group |  | Low-Responder group |  |
| --- | --- | --- | --- | --- |
|  | N=8 |  | N=6 |  |
|  | mean (SD) |  | mean (SD) |  |
| Age (years) | 69.62 (6.82) |  | 59.5 (13.47) | p=0.230 |
| Education | 12.25 (3.65) |  | 12 (2.82) | p=0.895 |
| Lesion volume (cm<sup>3</sup>) | 106.53 (110,60) |  | 228.98 (183,21) | p=0.230 |
| Lesion-testing delay (days) | 341.40 (340.59) |  | 816.33 (484.29) | **p=0.02** |
| Motor deficit (number of patients) | 4 |  | 3 |  |
| Visual field defect (number of patients) | 3 |  | 3 |  |
| **Neuropsychological results** |  |  |  |  |
| MMSE | 27.00 (1.69) |  | 26.83 (2.40) | p=0.881 |
| Recovery rate (%) | 36.29 (15.36) |  | -1.60 (22.18) | **p<0.01** |
|  | **Pre** | **Post** | **Pre** | **Post** |
| Neglect severity global score (%) | 32.38 (20,10) | 21.64 (16.69) | 32.18 (24.96) | 30.72 (19.53) |
| Letter cancellation (left omissions) | 10.25 (7.63) | 4.62 (6.09) | 7.83 (11.29) | 7.00 (9.63) |
| Gainotti drawing (max 6) | 4.50 (1.07) | 5.20 (1.00) | 4.33 (1.21) | 4.83 (1.33) |
| Reading task (max 116) | 8.50 (12.75) | 3.00 (3.82) | 16.00 (35.78) | 16.00 (37.24) |
| Line bisection (rightward deviation, mm) | 20.08 (22.43) | 8.35 (15.87) | 16.04 (28.89) | 10.23 (25.27) |
| Daisy drawing (max 3) | 2.00 (0.92) | 2.62 (1.06) | 1.33 (0.82) | 1.50 (0.84) |
| Landmark task ("left shorter" answers, max 12) | 8.00 (3.78) | 7.5 (3.74) | 8.00 (3.29) | 9.00 (3.29) |

For each assessment, patients were considered as showing neglect when they obtained pathological performance on at least three tests. A neglect severity score was computed as previously reported (Lunven et al., 2015), by using the percentage of left-sided omissions for letter cancellation, drawing and reading tasks, the percentage of rightwards deviation for line bisection, and the percentage of left lateralised errors for the landmark task. Taking into account only left-sided omissions can underestimate neglect in patients with very severe neglect, who do not reach the sheet midline and thus make also right-sided omissions. However, no patient in our sample showed this behaviour; all patients with pathological performance on letter cancellation showed a spatially lateralized deficit, with differences between left-sided and right-sided target omissions being greater than four targets) (see supplementary table 1)

### 2.3 Prism adaptation

Following a previously described procedure (Rossetti et al., 1998), patients were asked to wear prismatic goggles that shifted the visual field 10 degrees towards the right. While wearing prisms, patients were required to make, as quickly as possible, a series of approximately 100 pointing responses with the right hand towards left and right targets. The starting point of the hand was occluded to ensure optimal adaptation. The head of the patients was aligned with their body sagittal axis. Patients were eye-blinded and asked to point ten times straight-ahead with their right arm before and after the PA. Recovery scores were defined as the percentage of neglect improvement after PA, by using the following formula: (neglect severity score post-PA – neglect severity score pre-PA) / (neglect severity score pre-PA). The neglect severity score pre-PA was the score obtained for each subject at the second evaluation (two hours before PA). We adopted an arbitrary cutoff of 20% change, which is often used in clinical studies (Brunt et al., 2002; Cockcroft, 2010), to divide patients into two groups according to their recovery score. Patients with a recovery score above 20% were attributed to the “high-responder” group (N=8), whereas patients with a recovery score below 20% were considered as “low-responders” (N=6). This classification was in good agreement with the clinical assessment of neglect (see Table 1).

### 2.4 Imaging data acquisition and lesion analysis

#### 2.4.1 Data acquisition

An axial three-dimensional MPRAGE dataset covering the whole head was acquired for each participant (176 slices, voxel resolution = 1 × 1 × 1 mm, TE = 3 msec, TR = 2300 msec, flip angle = 9°) on a Siemens 3 T VERIO TIM system equipped with a 32-channel head coil.

Additionally, a total of 70 near-axial slices was acquired using an acquisition sequence which provided isotropic (2 × 2 × 2 mm) resolution and coverage of the whole head. The acquisition was peripherally-gated to the cardiac cycle with an echo time (TE) = 85 msec. We used a repetition time (TR) equivalent to 24 RR intervals (i.e., interval of time between two heart beat waves), assuming that spins would have fully relaxed before the repetition. At each slice location, six images were acquired with no diffusion gradient applied. Additionally, 60 diffusion-weighted images were acquired, in which gradient directions were uniformly distributed in space. The diffusion weighting was equal to a b-value of 1500 sec mm^-2^. One supplementary image with no diffusion gradient applied but with reversed phase-encode blips was collected. This provides us with a pair of images with no diffusion gradient applied with distortions going in opposite directions. From these pairs the susceptibility-induced off-resonance field was estimated using a method similar to that described in (Andersson, Skare, & Ashburner, 2003) and corrected on the whole diffusion weighted dataset using the tool TOPUP as implemented in FSL (Smith et al., 2004).

#### 2.4.2. Study of grey matter

Abnormal tissues were delineated on the T1-weighed images using MRIcroN for each patient (Rorden & Brett, 2000). T1-weighted images were normalized in the Montreal Neurological Institute (MNI) space (http://www.mni.mcgill.ca/) using rigid and elastic deformation tools provided in the software package Statistical Parametric Mapping 8 (SPM8, http://www.fil.ion.ucl.ac.uk/spm). Deformation was applied to the whole brain except for the voxels contained in the lesion mask, in order to avoid deformation of the lesioned tissue (Brett, Leff, Rorden, & Ashburner, 2001; Volle et al., 2008). Supplementary figure 1 displays the negative effect when normalization is done without the lesion mask. Finally, the lesion was carefully drawn in the MNI space by an expert neuropsychologist (ML) and checked by an expert neurologist (RM). Subsequently, lesions were overlapped for each of the high-responders and low-responders group.

#### 2.4.3. Cortical Thickness preprocessing

Cortical thickness was derived from the T1-weighted imaging dataset using a registration-based method (Diffeomorphic Registration based Cortical Thickness, DiReCT) (Das, Avants, Grossman, & Gee, 2009) optimised for patients with a brain lesion in BCBtoolkit (http://toolkit.bcblab.com) (Foulon et al., 2018). This approach has good scan-rescan repeatability and good neurobiological validity, as it can predict with a high statistical power the age and gender of the participants (Tustison et al., 2014). These data were then normalized to MNI space with the previous deformation estimated to normalize the T1-weighted imaging dataset.

#### 2.4.4. Study of white matter

We extracted brain using BET implemented in FSL and we corrected diffusion datasets simultaneously for motion and eddy current geometrical distortions by using ExploreDTI (http://www.exploredti.com) (Leemans & Jones, 2009). The tensor model was fitted to the data by using the Levenberg-Marquardt non-linear regression (Marquardt, 1963).

FA maps were further processed using Tract-Based-Spatial Statistics (TBSS) implemented in FSL (http://fsl.fmrib.ox.ac.uk/fsl/fslwiki/TBSS) (Smith et al., 2006). TBSS is the leading technique for voxel-wise DTI analysis, and it generally yields more reproducible results than alternative approaches (Bach et al., 2014). Comparing to voxel-based morphometry approach, TBSS may increase the statistical power of the analysis by focus comparison only on the white matter skeleton. TBSS also minimizes the chance that the results are driven by partial volume effects, because it avoids the need for smoothing the data. Moreover, TBSS requires minimal input from the user and has already been used with stroke patients, including neglect patients (Bozzali et al., 2012; Lunven et al., 2015; Vaessen, Saj, Lovblad, Gschwind, & Vuilleumier, 2016)

All patients’ FA data were aligned into a common space using the nonlinear registration tool FNIRT (Andersson 2007a, 2007b), which uses a b-spline representation of the registration warp fields (Rueckert 1999). In order to avoid deformation of the lesioned tissue during the registration we changed the original script implemented in FSL. For each patient we used her/his segmentation of lesion in the native space as mask during this step. Correct registration was visual checked for each patient (Lunven et al., 2015). Next, an average FA map was created and a skeleton map representing the centre of the white matter (FA > 0.2) common to all patients computed. Finally, the registered FA maps were projected into the skeleton. The nonlinear warps and skeleton projection were be applied to MD, RD and AD images.

Secondly, we used the Disconnectome maps of the BCBtoolkit to further characterise the white matter tracts passing through the voxels showing a significant difference. Diffusion weighted imaging datasets were obtained from 10 healthy controls (Rojkova et al., 2015). Tractography was estimated as described in (Thiebaut de Schotten, Dell’Acqua, et al., 2011). Patients’ lesions in the MNI 152 space are registered to each control native space using affine and diffeomorphic deformations (Avants et al., 2011; Klein et al., 2009) and subsequently used as seed for the tractography in Trackvis (http://www.trackvis.org/). Tractography from the ROIs were transformed in visitation maps (Thiebaut de Schotten, ffytche, et al., 2011), binarized and brought to MNI 152 using the inverse of precedent deformations. Finally, we produced a percentage overlap map by summing at each point in MNI space the normalized visitation map of each healthy subject. Hence, in the resulting disconnectome map, the value in each voxel take into account the interindividual variability of tract reconstructions in controls, and indicate a probability of disconnection from 0 to 100% for a given brain ROI (Thiebaut de Schotten et al., 2015).

#### 2.4.5. Statistical analysis

We statistically compared FA and cortical thickness values between low-responder and high-responder patients. Visual field defects and lesion volume were included as co-variables of non-interest. Lesioned voxels were excluded from these analyses. We used the FSL function “Randomise”, with 5,000 random permutation tests and a Threshold-Free Cluster Enhancement option. The method takes a raw statistic image and produces an output image in which the voxel-wise values represent the amount of cluster-like local spatial support. This method improves sensitivity and is more interpretable output than cluster-based thresholding (Smith & Nichols, 2009). Results were adjusted for FWE corrections for multiple comparisons and thresholded at p<0.05. In addition, we applied the same analyses to explore potential between-group differences in mean diffusivity, axial diffusivity and radial diffusion.

To assess the dimensional relationships between FA, MD, RD, AD, cortical thickness and prism-induced changes in performance, we conducted a regression analysis including all the 14 patients. Lesion volume and visual field defect were included as continuous co-variables of non-interest. We again used the FSL function “Randomise”, with 5,000 random permutation tests and a Threshold-Free Cluster Enhancement option. The method takes a raw statistic image and produces an output image in which the voxel-wise values represent the amount of cluster-like local spatial support. This method improves sensitivity and is more interpretable output than cluster-based thresholding (Smith & Nichols, 2009). Results were adjusted for FWE corrections for multiple comparisons and thresholded at p<0.05.

In a final step, we adopted a ROI-based approach to test FA and cortical thickness values differences between controls and the two subgroups of patients. Healthy participants could not be included in the TBSS analysis model because of the regression for the lesion size included in the model, which is not adapted to a control group without lesion.

For each statistical analysis, effect sizes were calculated using G*power (http://www.gpower.hhu.de/en.html).

## 3. Results

### 3.1. Behavioural results

PA induced a sensorimotor adaptation in all patients, who deviated rightward before PA (6.47 degrees; SD, 3.51), but leftward after PA (−1.95 degrees; SD, 2.83; Z=-3.234, p=0.001). This adaptation was defined when leftward deviation was at least 3 cm (Luauté et al., 2006). There was no correlation between neglect recovery scores and PA effect on the straight-ahead task, consistent with previous results (Pisella, Rode, Farnè, Boisson, & Rossetti, 2002).

Our patient sample was relatively small (n<20), thus we performed non-parametric statistical tests on demographic and behavioural data. Patients demonstrated a relatively stable level of performance before PA, with at least 3 pathological tests for the two pre-tests; neglect scores obtained one day and two hours before PA were similar (first session: mean score 30.83, SD 21,39; second session: mean score 32.29, SD 21.38; Wilcoxon Signed-Ranks Test, Z=-1.73, p=0.084). The mean percentage of performance change for each patient [calculated as (neglect severity score “2 hours before PA” – neglect severity score “the day before PA”) / (neglect severity score “the day before PA”)] was 5.43% (SD: 13.20, min −15.46, max 30.18). Patient 7 and 13 obtained worse scores at the second evaluation than at the first (by respectively 30.18% and 20.38%, supplementary table 1). Patient 7 was considered as high-responder because she improved more than 20% between the second pre-test and the post-PA test. Even when we compared the evolution of performance between the first pre-test and the post-PA test, we found an improvement of 48.33% in this patient. Patient 13 was included in the low-responder group, because his score did not change between the second pre-test and the post-PA test; he only improved by less than 20% (19.2%) between the first pre-test and the post-PA test.

High-responder and low-responder patients had similar age (Mann-Whitney U=34; p=0.23), with high-responders being numerically older (mean, 69.62 years, SD 6.82) than low-responders (59.50 years, SD 13.47). Low-responders had numerically larger lesion volumes, but the difference did not reach significance (U=14; p=0.23). Low- and high-responders had similar neglect severity scores before PA (U=28; p=0.66). However, high-responders had a shorter delay from stroke (mean number of days from stroke, 341.38; SD, 340.58; range 107-1119) than low-responders (mean, 816.33 days; SD, 484; range 319-1497; U=42, p=0.02). Homonymous hemianopia was present in three low-responders and in four high-responders.

By definition, high-responder patients demonstrated a statistically reliable neglect improvement after PA (Wilcoxon Signed-Ranks Test, Z=-2.52; p=0.012), while this was not the case for the low-responder subgroup (Z=-0.67; p=0.50). In particular, high-responders detected more left-sided items in letter cancellation (Z=-2.39, p=0.017), drew more left-sided items in landscape copy (Z=-2.06, p=0.039) and in daisy drawing (Z=-2.24; p=0.025), and had a more symmetrical performance when bisecting lines (Z=-2.240; p=0.025). However, PA did not significantly impact performance on tasks with minimal motor component, such as the reading task (Z=-1.604; p=0.11) and the Landmark task (Z=-0.365; p=0.715).

### 3.2. Lesion analysis

The maximum lesion overlap was centred in the right fronto-parietal white matter in both patient subgroups (Figure 1.1). In low-responders, damage also involved the supramarginal and angular gyri of the inferior parietal lobule. Lesion subtraction (Figure 1.2) between low and high responders mostly revealed parietal regions, including the inferior parietal lobule, the postcentral gyrus, but also frontal regions such as the precentral gyrus, the superior frontal lobe, the orbito-frontal cortex, and the fronto-parietal white matter.

**Figure 1:**
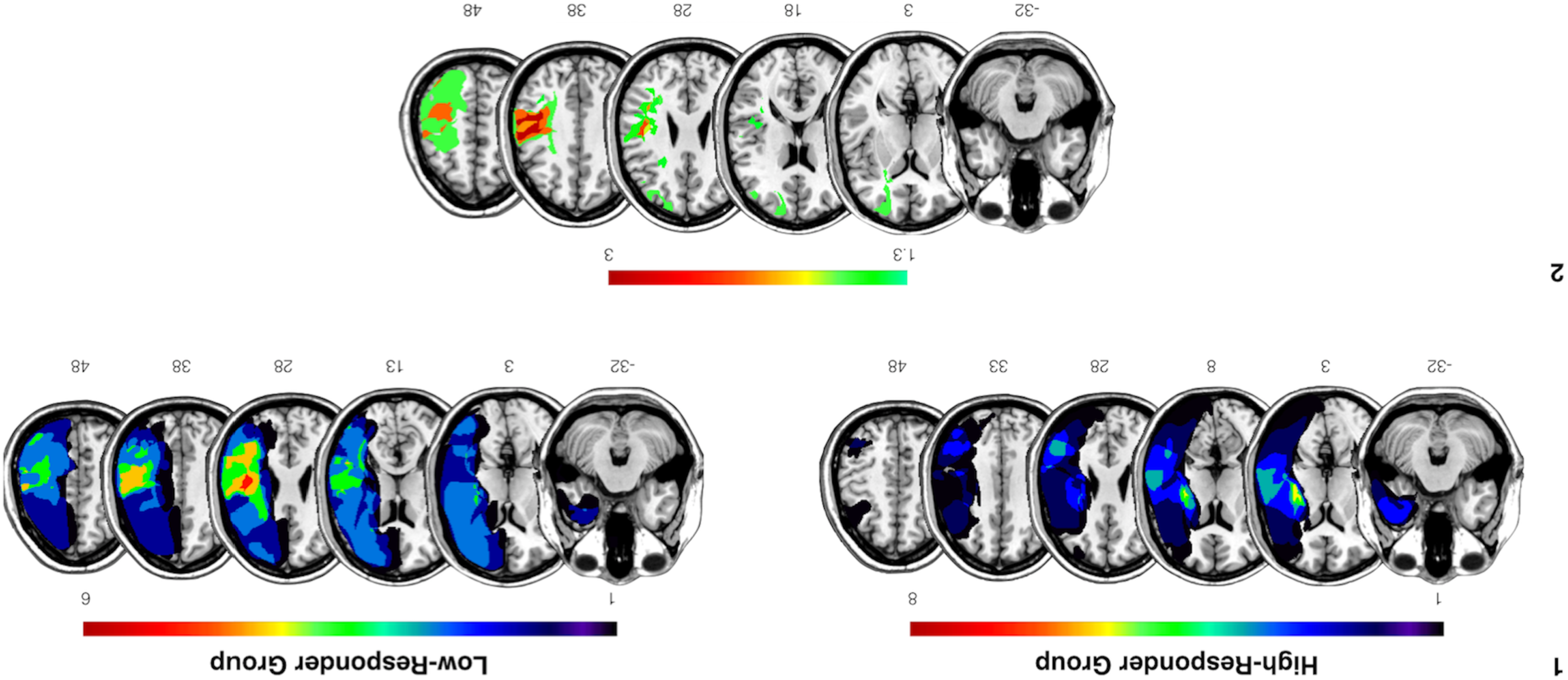
Patients’ grey matter lesion anatomy. (1) Overlap of the lesions for all the included right brain-damaged patients for each group. (2) Results of the subtraction of the probability map of the high-recovery group from the probability map of the low-recovery group. Maps show the Z-statistics rank order statistics. All peaks are significant at p<0.05 level.

**Figure 2:**
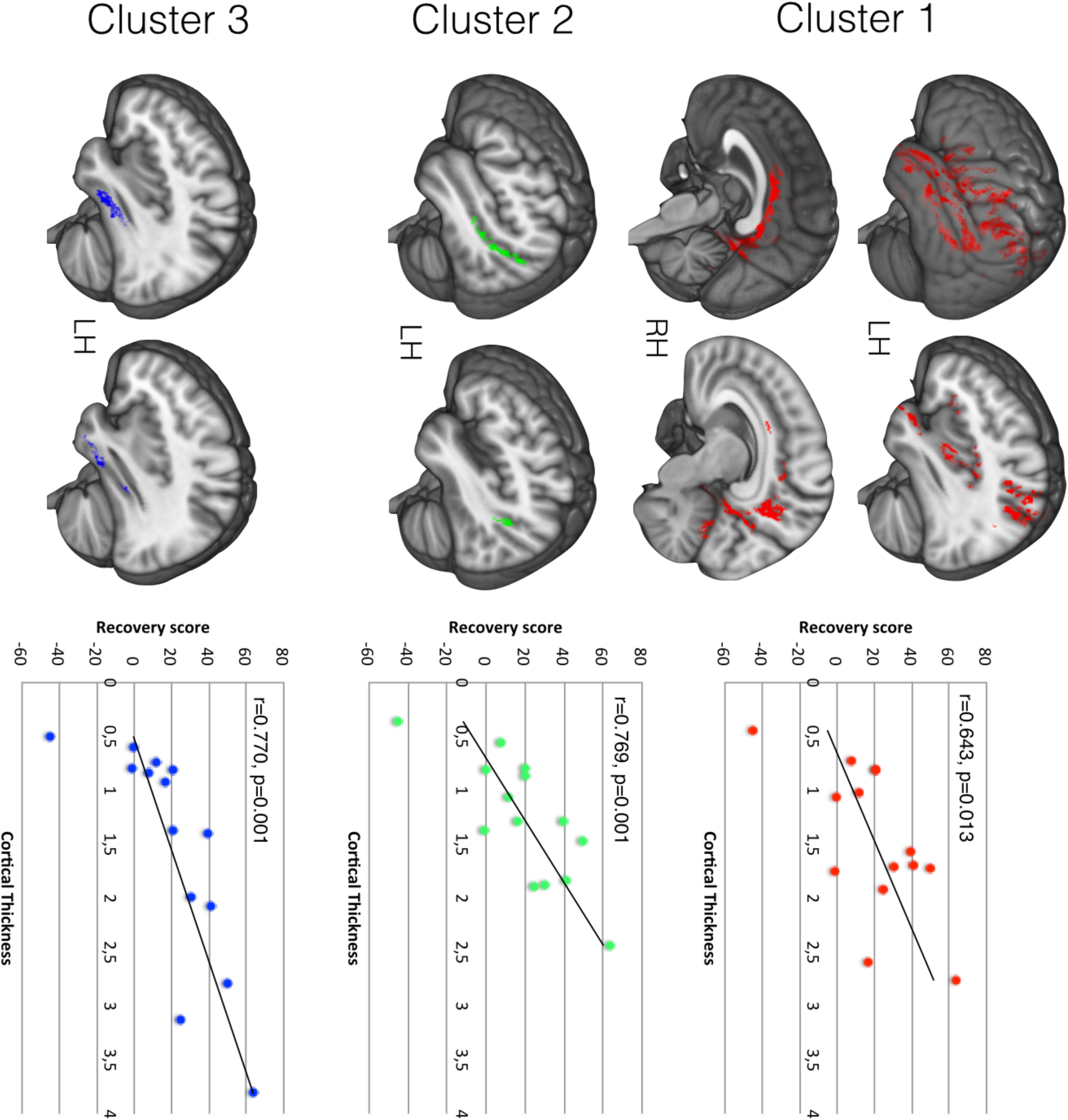
Cortical thickness analysis. Relationship between cortical thickness and recovery rate score represented by the four clusters. Results are represented at p<0.05 and corrected for multiple comparisons. LH, left hemisphere; RH, right hemisphere.

### 3.3. Cortical thickness

A regression analysis between cortical thickness and recovery scores revealed three significant clusters in the left, undamaged hemisphere (Figure 2 and Table 2). The first cluster (p=0.037) involved the perisylvian region, including the surface of the inferior parietal lobule, the three temporal gyri, somatosensory areas and the prefrontal cortex, with a medium effect size (0.26). The second cluster (p=0.048) was located in the posterior part of the superior temporal sulcus. The third cluster (p=0.049) was situated in the anterior portion of the left inferior temporal sulcus. Two of the three clusters had large effect sizes (>0.60, Cohen, 1988).

**Table 2:**
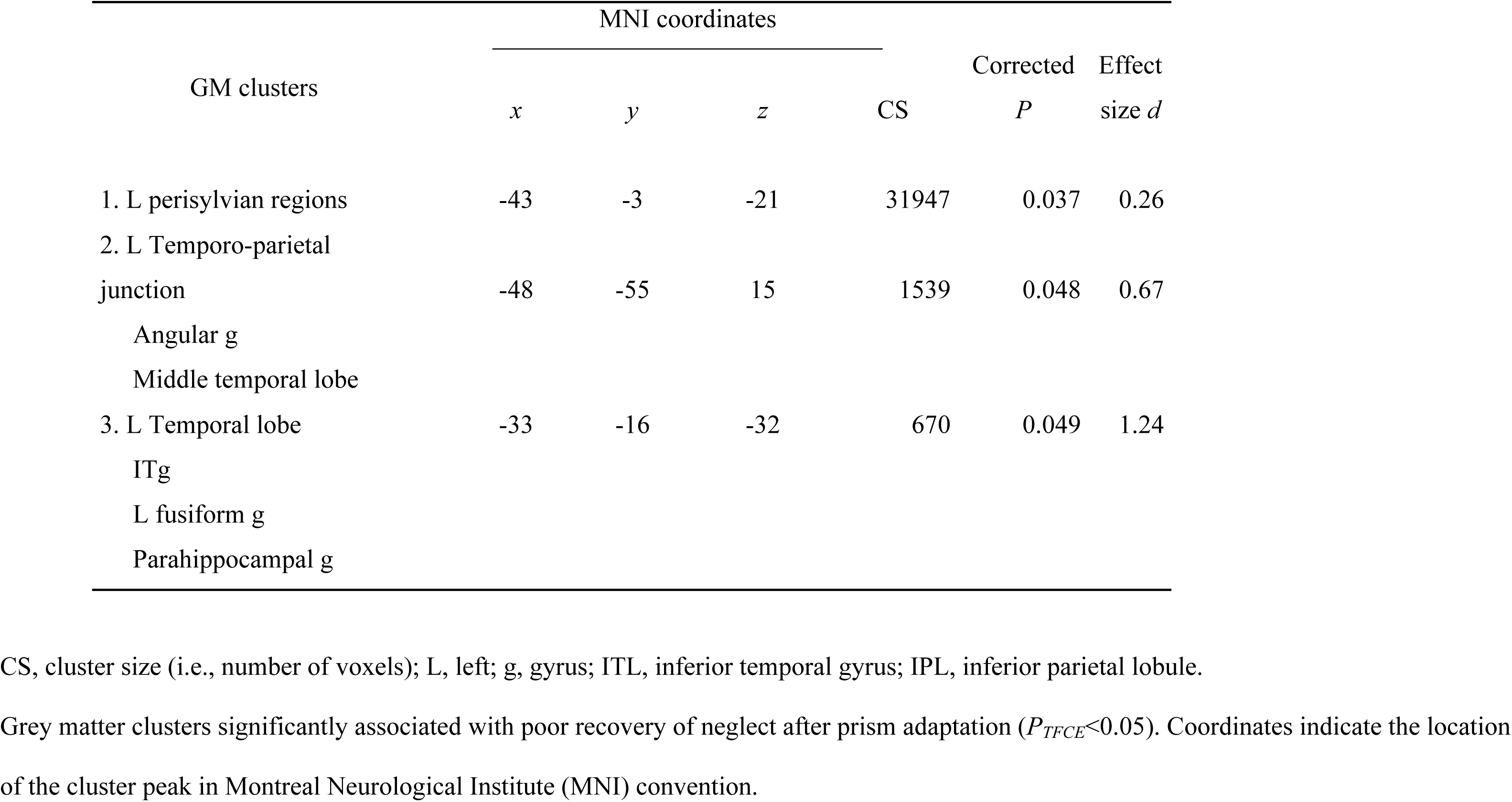
Grey matter clusters showing decreased cortical thickness associated with poor recovery after PA therapy CS, cluster size (i.e., number of voxels); L, left; g, gyrus; ITL, inferior temporal gyrus; IPL, inferior parietal lobule. Grey matter clusters significantly associated with poor recovery of neglect after prism adaptation (*P*_*TFCE*_<0.05). Coordinates indicate the location of the cluster peak in Montreal Neurological Institute (MNI) convention.

Controls and high responders had similar values of cortical thickness; low responders had instead lower values in cluster 3 (mean, 0.73) compared with high-responders (2.17) (t=3.9751, df=12, Benjamini-Hochberg-corrected p<0.01), and with controls (mean=2.52) (t=9.48; df=14; Benjamini-Hochberg-corrected p<0.01) (Supplementary Figure 2). There were statistical trends for differences in the same direction in cluster 2 between low-responders and high-responders (means, 0.91 and 1.55, respectively; t=2.51; df=12; Benjamini-Hochberg-corrected p = 0.082), and between low-responders and controls (controls’ mean, 1.47; t=2.88; df=14; Benjamini-Hochberg-corrected p=0.055).

For each cluster, we conducted a linear regression with the cortical thickness values, the recovery score and the number of days post stroke. The linear regression models did not reveal any association between values of cortical thickness, the recovery score and the number of days post stroke.

### 3.4. White matter analyses

TBSS showed that low-responders had decreased FA as compared with high-responders in three clusters, all with large effect sizes (>0.80; Figure 3, Table 3). The first cluster (p = 0.019) was located in the body of the corpus callosum. The second cluster (p = 0.049) encompassed the genu of the corpus callosum. The third cluster (p = 0.049) was localized in a portion of the genu of the corpus callosum within the left, undamaged hemisphere.

**Table 3:** White matter clusters showing decreased fractional anisotropy FA (*P*_*TFCE*_<0.05) in the low-responder group as compared with the high-responder group CS, cluster size (i.e., number of voxels); CC, corpus callosum; L, left. Coordinates indicate the location of the cluster peak in Montreal Neurological Institute (MNI) convention.

| WM clusters | MNI coordinates | | | CS | Corrected $P$ | Effect size $d$ |
| --- | --- | --- | --- | --- | --- | --- |
| | $x$ | $y$ | $z$ | | | |
| 1. CC body | 2 | -20 | 24 | 1533 | 0.019 | 1.57 |
| 2. CC genu | 0 | 24 | 0 | 47 | 0.049 | 1 |
| 3. L CC genu | -12 | 31 | -2 | 9 | 0.049 | 0.85 |

CS, cluster size (i.e., number of voxels); CC, corpus callosum; L, left.

**Figure 3:**
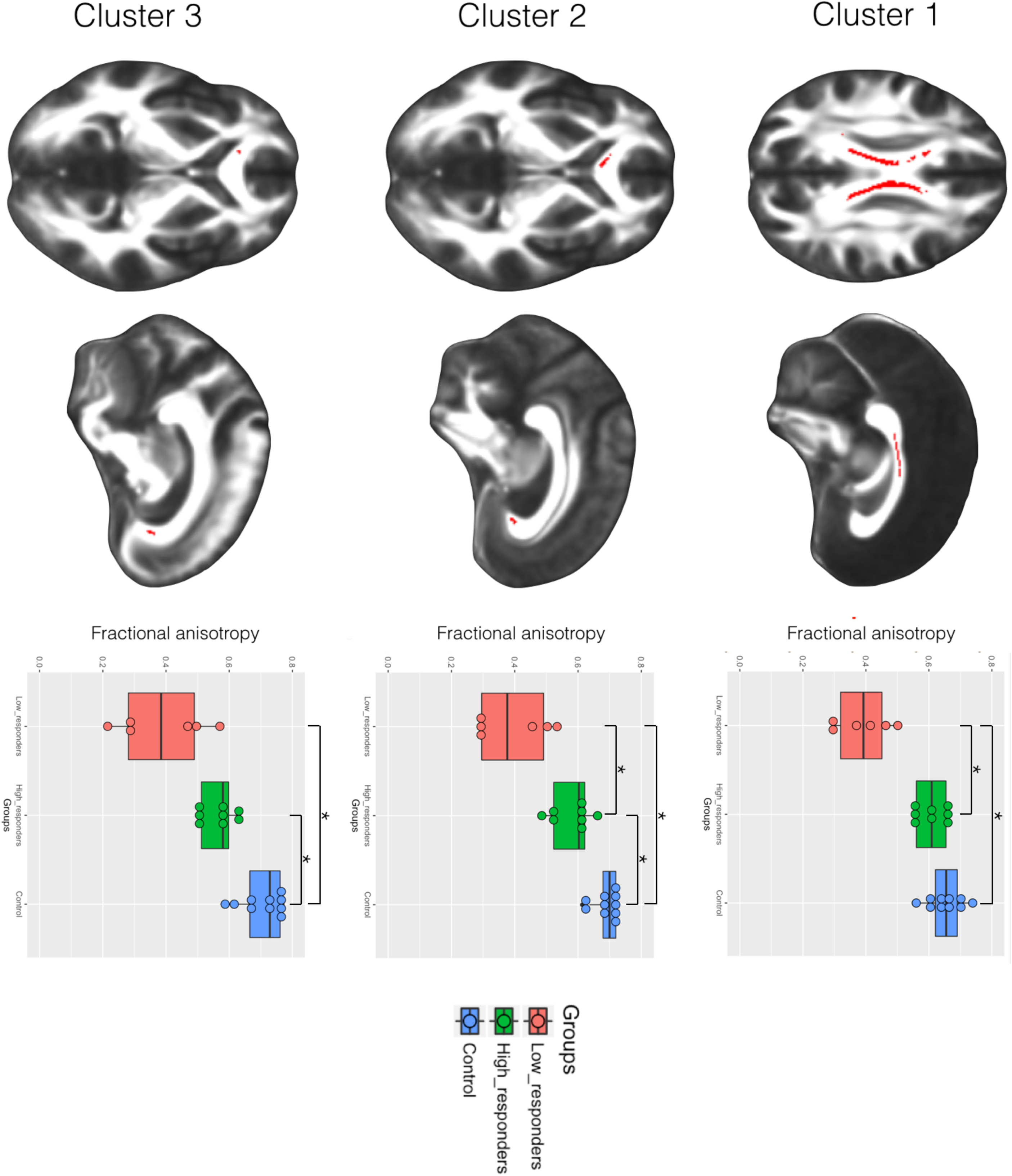
White matter analysis. Results of TBSS between-groups comparisons represented by the three clusters. FA differences (p<0.05) are represented in red.

We used BCBtoolkit to visualize the white matter tracts passing through the voxels showing a significant difference between the patient subgroups. The analysis revealed that the fibres intersecting the first cluster mostly connected sensorimotor areas (Figure 4). Additional regions in the temporal lobe (middle and superior gyri), and in the parietal lobe (supramarginal gyrus, precuneus and superior parietal lobule) were also involved. The second and the third clusters mostly connected prefrontal areas. TBSS analysis did not uncover group differences in mean, axial and radial diffusivity. Regression analyses between FA, MD or AD values and percentage of improvement after PA did not disclose any significant correlations.

**Figure 4:**
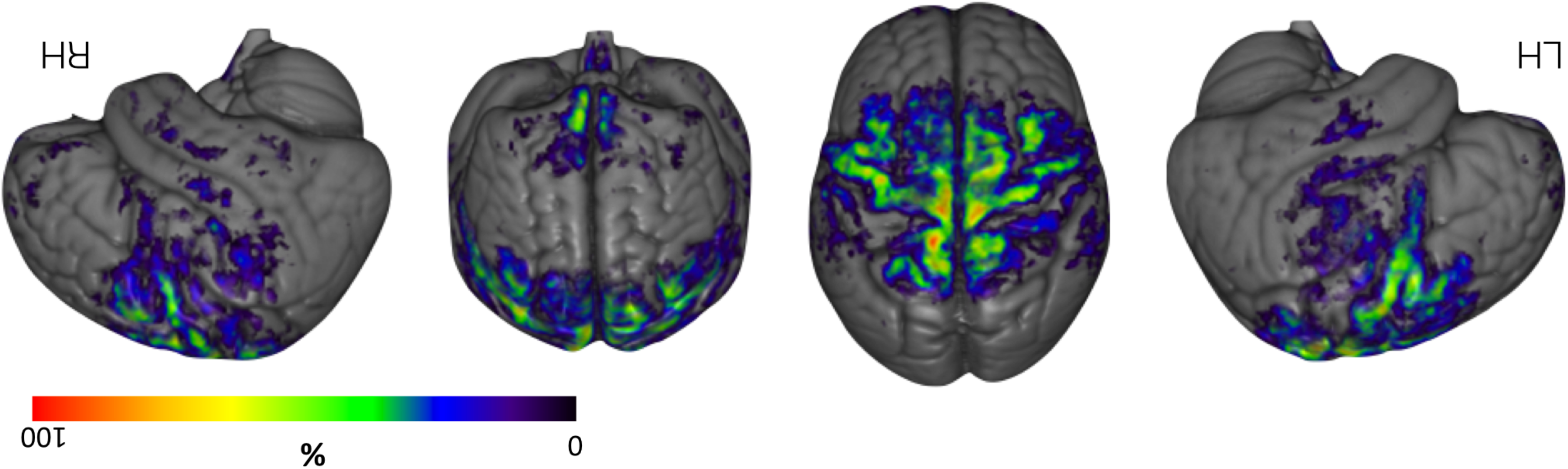
Percentage maps in healthy subjects (BCB toolkit: http://www.brainconnectivitybehaviour.eu) showing the cortical projections of the voxels identified by TBSS analysis in low recovery patients compared to high-recovery patients. LH, left hemisphere; RH, right hemisphere.

A region of interest approach based on the clusters identified by TBSS analysis in patients was employed to extract FA values from the ten age- and education-matched healthy participants. Benjamini-Hochberg-corrected comparisons showed similar FA values in high-responders and controls in cluster 1 (p=0.30). Low-responders had lower FA values than both controls and high-responders in cluster 1 (all p<0.01) and 2 (all p<0.01). High- and low-responders had lower FA values than controls in cluster 3 (p<0.01), with a tendency for lower FA values for low-responders than for high-responders (p= 0.081) (Figure 3).

For each ROI representing one cluster, Spearman’s rank correlation test was performed to determine the strength of the relationship between FA and cortical thickness values. Cortical thickness in clusters 2 and 3 correlated with the 3 clusters of FA values (Table 4), thus suggesting a link between decreased cortical thickness and white matter degeneration. All FA and Cortical thickness clusters correlated with the delay post-stroke (in days, table 5).

**Table 4:** Spearman correlations between the three FA clusters identified in the TBSS analysis and the three cluster identified in the cortical thickness study

|  | FA Cluster 1 | FA Cluster 2 | FA Cluster 3 | CT Cluster 1 | CT Cluster 2 | CT Cluster 3 |
| --- | --- | --- | --- | --- | --- | --- |
| FA Cluster 1 |  |  |  |  |  |  |
| FA Cluster 2 | **0.855\*\*** |  |  |  |  |  |
| FA Cluster 3 | **0.780\*\*** | **0.864\*\*** |  |  |  |  |
| CT Cluster 1 | 0.253 | 0.358 | 0.437 |  |  |  |
| CT Cluster 2 | **0.582\*** | **0.692\*\*** | **0.705\*\*** | **0.873\*\*** |  |  |
| CT Cluster 3 | **0.701\*\*** | **0.767\*\*** | **0.771\*\*** | **0.648\*\*** | **0.798\*\*** |  |

\*\* p<0.01; \* p<0.05, FA Fractional Anisotropy, CT Cortical Thickness,

**Table 5:** Spearman correlations between imaging clusters and delay post stroke

|  | Cluster 1<br>FA | Cluster 2<br>FA | Cluster 3<br>FA | Cluster 1<br>CT | Cluster 2<br>CT | Cluster 3<br>CT |
| --- | --- | --- | --- | --- | --- | --- |
| Correlation coefficient with delay post stroke | -0.71** | -0.58* | -0.59* | -0.49* | -0.65** | -0.62** |

\*\* p<0.01; \* p<0.05, FA Fractional Anisotropy, CT Cortical Thickness,

## 4. Discussion

In this study, we followed up a group of patients with chronic visual neglect, before and after prism adaptation. The time elapsed from stroke resulted to be shorter in high-responders than in low-responders, which suggests the opportunity to administer PA therapy as soon as possible after the stroke. We used advanced neuroimaging techniques to identify the anatomical predictors of neglect patients’ response to PA. First, cortical thickness analyses indicated a significant contribution of the left, undamaged hemisphere. Second, diffusion weighted imaging demonstrated an important role for inter-hemispheric connections in PA efficiency.

In the left, healthy hemisphere, cortical thickness analyses revealed a significant involvement of the temporo-parietal areas in the PA cognitive effect. These regions match areas contributing to right visual neglect after a stroke in the left hemisphere (Beume et al., 2016), and mirror classical areas reported as involved in left visual neglect after a stroke in the right hemisphere (Corbetta & Shulman, 2011). This may suggest that the left temporo-parietal junction is able to compensate for left visual neglect (Paolo Bartolomeo & Thiebaut de Schotten, 2016). Consistent with this hypothesis, previous studies reported an involvement of the left hemisphere in the recovery of neglect after PA therapy (Levin, Kleim, & Wolf, 2009; Luauté et al., 2006; Saj et al., 2013). In healthy individuals, rightward PA increased BOLD response in the left inferior parietal cortex (Tissieres, Fornari, Clarke, & Crottaz-Herbette, 2017). Our cortical thickness and diffusion results suggest an implication of left prefrontal cortex in prism-induced neglect compensation, consistent with the observation that effective PA in neglect is facilitated by TMS-based disinhibition of the left motor cortex (O’Shea et al., 2017). Altogether, these findings fit well with the fronto-parietal organization of attention networks in the primate brain (Lunven & Bartolomeo, 2017).

Two possibilities can explain the difference in cortical thickness in the contralesional hemisphere. Some individual brains might have a better predisposition to recover than others (e.g., thanks to a thicker left temporo-parietal junction). This possibility would be consistent with the analogous results obtained in the domain of recovery from aphasia, which appears to be promoted by stronger language structures in the right hemisphere (Forkel et al., 2014; Xing et al., 2016). An alternative hypothesis might be that diaschisis and disconnection may have induced secondary atrophy in remote cortices (Cheng et al., 2015; Foulon et al., 2017). This would be consistent with previous reports suggesting that bilateral diffuse tissue loss may occur in individuals with chronic stroke (Gauthier, Taub, Mark, Barghi, & Uswatte, 2012; Kraemer et al., 2004). Thus, premorbid atrophy in left temporo-parietal areas might hamper cognitive effects related to PA after a stroke in the right hemisphere (see Levine, Warach, Benowitz, & Calvanio, 1986). Future longitudinal studies comparing acute to chronic stages of vascular strokes may adjudicate between these possibilities.

Our results also showed that low-responder patients had a selective decrease of callosal FA in the body and genu of the corpus callosum, which mainly connect sensorimotor and prefrontal areas (Catani & Thiebaut de Schotten 2012). Martín-Arévalo *et al.* (2016) showed that leftward PA alters transcallosal motor inhibition between the two motor cortices in the healthy brain, thus confirming the possibility that PA modulates inter-hemispheric interactions. This result suggests that PA improves left neglect by allowing left-hemisphere attentional networks to access information processed by the lesioned right hemisphere, through the callosal body and genu. An implication of prefrontal circuits in neglect compensation has been hypothesized on the basis of behavioural evidence (Bartolomeo, 1997; Bartolomeo, 2000), and recently confirmed by electrophysiological measures (Takamura et al., 2016). Also consistent with our results, cathodal transcranial direct current stimulation applied over the left posterior parietal cortex interfered with the beneficial effect of PA on neglect (Làdavas et al., 2015).

## 5. Limitations of the study

We were able to include 14 chronic neglect patients for follow-up before and after PA; the design of the study required this sample to be further subdivided in two subgroups (high- and low-responders). The limited number of inclusions is representative of the difficulties of organizing a longitudinal study of patients with stable neuropsychological deficits, with the associated need of administering lengthy MRI sequences and repeated cognitive assessments. In the present study, five additional neglect patients had to be excluded because they did not tolerate the 1-hour-long neuroimaging procedure. Despite this limitation, however, the sample size was sufficient to obtain significant results with TBSS, which is a notoriously conservative technique. Also, a power analysis indicated medium to large effect sizes for the left temporo-parietal junction, temporal lobe and white matter findings. Finally, the present anatomical evidence is consistent with previous studies, showing that lesions to the right hemisphere posterior parietal cortex and the superior frontal gyrus (Rousseaux et al., 2006), the infero-posterior parietal cortex (Luauté et al., 2006), the intra-parietal sulcus, the white matter around the inferior parietal lobule, the middle frontal gyrus (Sarri et al., 2008), the occipital lobe (Serino et al., 2006) and the right cerebellum [patient 5 in Saj et al., 2013)] hampered PA efficiency. Thus, our patient population appears to be representative of the general lesional patterns in chronic neglect patients.

## 6. Conclusions and perspectives

Our results confirm the importance for recovery from neglect of the left, healthy hemisphere (Bartolomeo & Thiebaut de Schotten, 2016), as well as of efficient inter-hemispheric connections (Lunven et al., 2015). Specifically, left hemisphere regions homologues to the fronto-parietal attention networks in the right hemisphere (Corbetta & Shulman, 2002), as well as callosal pathways passing through the body and the genu, appear to be critical for compensation of neglect signs after a single PA session. It remains to be seen whether or not similar anatomical constraints apply to more prolonged rehabilitation programmes using PA (Frassinetti, Angeli, Meneghello, Avanzi, & Làdavas, 2002). The identification of anatomical predictors of response to different, labour-intensive rehabilitation strategies may ultimately enable clinicians to select and tailor the appropriate treatment to the specific needs of the individual patient. Even in chronic patients at more than 3 months from the stroke, early administration of PA could be important to improve patients’ response. Also, the relationship between recovery and contralesional cortical thickness suggests potential targets for future therapeutic interventions. Finally, we note that within a few months, a brain lesion affects cortical macro- and microstructures remotely, significantly reducing cortical thickness in distant regions connected to the lesion (Schaechter, 2006; Foulon et al. 2018). Consistent with these findings, our results indicate that the number of days post-stroke and the cortical thickness have a collinear contribution to prism adaptation effect. This suggests that prism adaptation should be administered as early as possible after stroke, to optimize the chances for recovery before long term disconnection effect impact the contralesional cortex. Future studies should investigate the complex tripartite relationship between cortical thickness change and PA reeducation effect by using following up patients not only behaviorally, but also with serial neuroimaging assessments.

### Study funding

The research leading to these results has received funding from a CIFRE PhD fellowship to ML, from the ANR “Phenotypes” ANR-13-JSV4-0001-01 to MTS and from the programs “Investissements d’Avenir” ANR-10-IAIHU-06, ANR-10-LABX-0087 IEC and ANR-10-IDEX-0001-002-PSL*.

## Acknowledgments

We would like to thank Cécile Coste, Fanny Lagneau, Karynne Moreau, Emilie Monnot and Marika Urbanski for patient referral. The research leading to these results has received funding from a CIFRE PhD fellowship to ML, from the ANR “Phenotypes” ANR-13-JSV4-0001-01 to MTS and from the programs “Investissements d’Avenir” ANR-10-IAIHU-06, ANR-10-LABX-0087 IEC and ANR-10-IDEX-0001-002-PSL*.

## Figure legends

**Supplementary figure 1:**
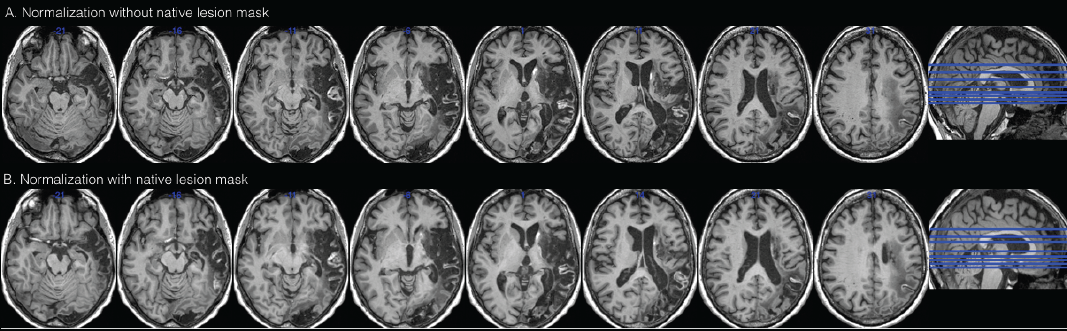
Negative effect on brain registration without including the lesion mask.

**Supplementary figure 2:**
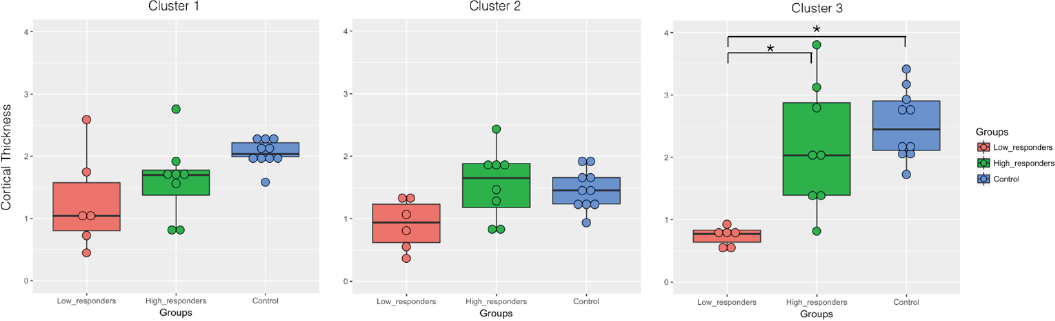
Comparison of cortical thickness values between the three groups for each cluster identified in the “randomise” analysis.

**Supplementary table 1:**
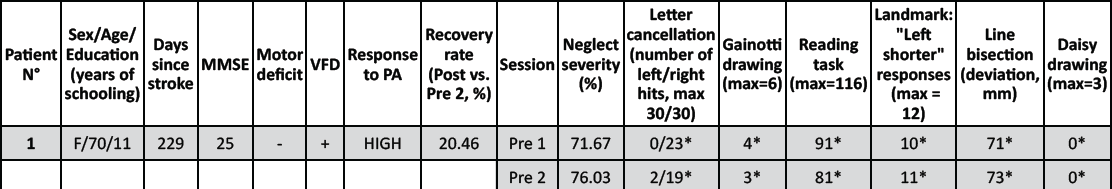
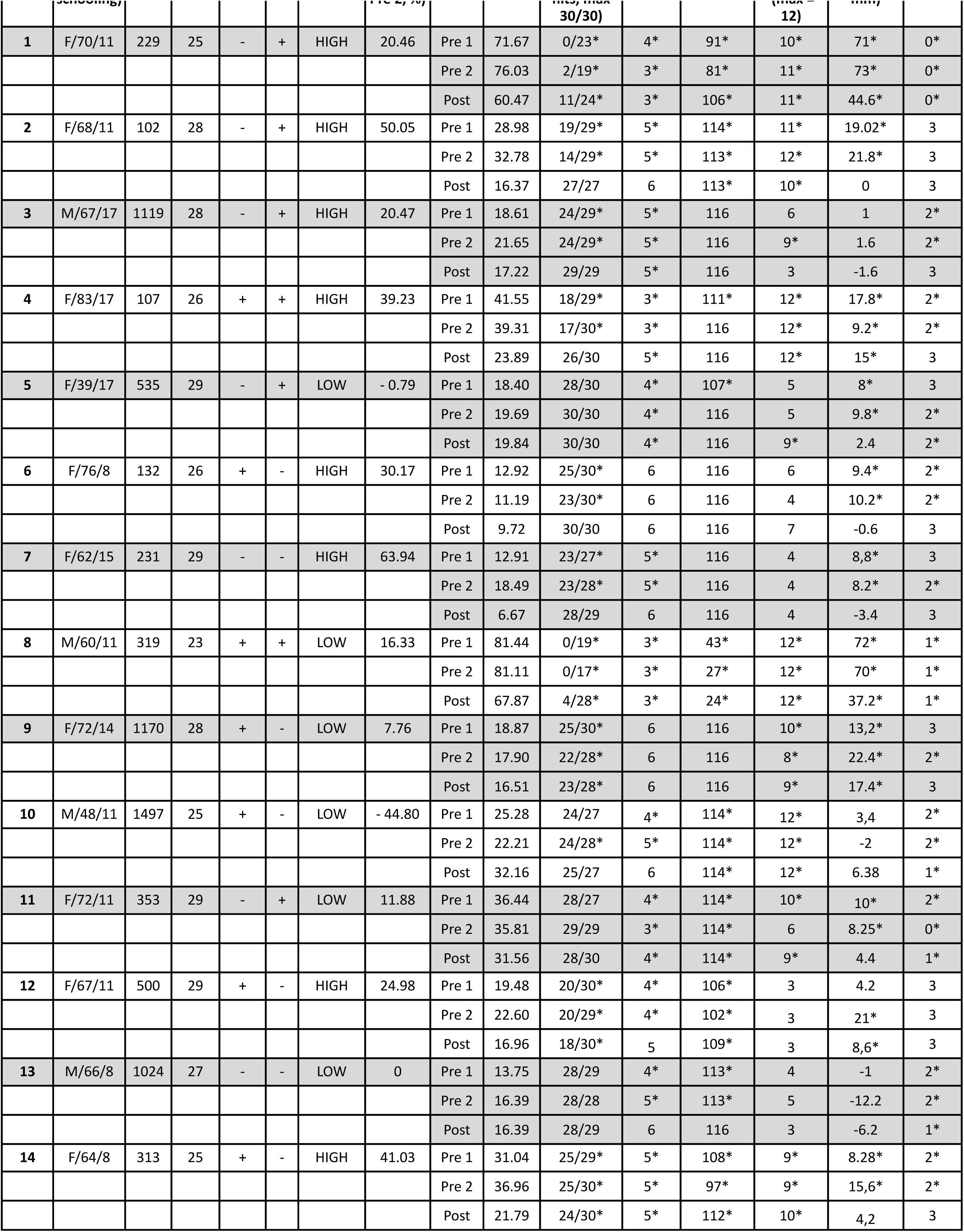
Demographic and clinical data of patients

